# Patterns and correlates of invasive alien plant richness in China’s abandoned croplands: a literature-derived synthesis

**DOI:** 10.64898/2026.07.23.740411

**Authors:** Xinlei Guo, Shuihua Wang, Yiming Huang, Ziyue Hong, John Moraros

## Abstract

Cropland abandonment is a widespread socio-ecological transition that can generate heterogeneous ecological outcomes, including opportunities for vegetation recovery as well as the accumulation of invasive alien plants (IAPs). Understanding broad-scale correlates of IAP richness in abandoned croplands is challenging because published studies differ in sampling design, spatial extent, and reporting detail. To provide an exploratory national-scale synthesis, we compiled literature-derived occurrence records of 57 IAP species reported from abandoned croplands across China and examined study-level recorded richness using generalized linear mixed models (GLMMs) with province as a random effect. Fourteen socio-economic and environmental predictors were screened for multicollinearity and standardized, and the number of Higher Education Institutions (HEIs) was included as a proxy for potential sampling-effort variation. Asteraceae accounted for 43.9% of recorded species, and more than 70% of taxa were classified as high-risk invaders (Levels 1-2). Recorded richness exhibited a pronounced southeast-northwest gradient, with higher values in economically developed coastal provinces. In the exploratory GLMM, regional GDP (IRR = 1.67, p < 0.05) and mean annual temperature (IRR = 1.49, p < 0.05) were positively associated with recorded richness, whereas other environmental, accessibility, soil, and sampling-effort variables showed no detectable associations. Model diagnostics indicated no evidence of overdispersion, zero inflation, or residual spatial autocorrelation. Given the heterogeneous nature of literature-derived data and the small number of study units, these results should be interpreted as broad-scale correlational patterns rather than causal drivers. The findings highlight regional socio-economic context and climatic conditions as key correlates of recorded IAP richness in abandoned croplands and provide an initial baseline for understanding invasion patterns in post-agricultural landscapes undergoing rapid land-use transitions..

## 1. Introduction

Land-use and land-cover change (LULCC) is an important component of global environmental change in the Anthropocene^1^. Among contemporary land-use transitions, cropland abandonment (CA) has become increasingly widespread, driven by interacting socioeconomic and environmental processes, including rural depopulation, labor migration, agricultural marginalization, and land degradation^2–6^. Globally, more than 472 million hectares of cropland have been abandoned, with over 100 million hectares affected between 1992 and 2020^7,8^. CA is frequently linked in the literature to vegetation recovery and carbon accumulation^9–11^, it also represents a novel disturbance regime that may facilitate biological invasions, particularly by invasive alien plants (IAPs)^12–15^.

From an invasion ecology perspective, cropland abandonment constitutes a transitional socio-ecological system in which the cessation of direct agricultural management coincides with persistent landscape connectivity and residual human influence. Such conditions may simultaneously enhance propagule pressure through surrounding anthropogenic matrices while altering environmental filtering processes that regulate establishment success. However, the relative roles of socio-economic drivers (e.g., human mobility, economic development) versus environmental constraints (e.g., climate, soil, and topography) in structuring invasion patterns in abandoned croplands remain poorly quantified at broad spatial scales^16^.

Invasive alien plants are widely recognized as an important component of global change biology due to their documented impacts on biodiversity, ecosystem functioning, agricultural systems, and landscape dynamics^17–23^. Conceptual frameworks in invasion ecology highlight the relevance of propagule availability, environmental constraints, and biotic interactions for understanding where alien species are more frequently observed^24–25^. In human-dominated landscapes, propagule availability is often linked to economic activity, transportation networks, and land - use intensity, whereas environmental constraints are associated with climatic and edaphic conditions that vary across regions^16,26^. Although these concepts provide a useful interpretive context, broad-scale empirical assessments of how such correlates align with recorded IAP richness in abandoned croplands remain limited, particularly when multiple predictor classes and potential sampling biases must be considered simultaneously.

In China, rapid economic development, expanding transportation networks, and extensive land - use transitions have coincided with increasing reports of alien plant occurrences across multiple regions^27–30^. At the same time, the country spans pronounced climatic and environmental gradients that may contribute to regional differences in where invasive alien plants are more frequently documented. Several widely recognized invasive species-including *Alternanthera philoxeroides*, *Spartina alterniflora*, *Ipomoea cairica*, *Solidago canadensis*, and *Ambrosia trifida*, have been recorded across diverse landscapes^16^. Despite this growing body of research, most studies of abandoned croplands have focused on local ecological processes such as soil seed banks, species interactions, or soil properties^31–33^. Consequently, national-scale evidence on how socio-economic and environmental correlates align with recorded IAP richness in abandoned croplands remains limited. Few studies have jointly considered multiple predictor classes while addressing multicollinearity and potential spatial or sampling biases inherent in literature-derived datasets^34–36^.

To address these gaps, the present study synthesizes literature -derived occurrence records of invasive alien plants reported in studies of abandoned croplands across China. Specifically, we aim to: (1) describe the taxonomic composition and invasion - risk categories of recorded alien plant assemblages; (2) characterize broad -scale spatial variation in recorded IAP richness; and (3) explore how socio-economic, climatic, edaphic, and topographic variables align with study-level recorded richness using generalized linear mixed models (GLMMs), while accounting for spatial clustering and potential sampling-effort variation. Given the heterogeneous nature of literature-derived data and the limited number of study units, the analysis is exploratory in scope and the results reflect broad-scale correlational patterns rather than causal relationships. This national-scale synthesis provides an initial macroecological overview of regional correlates of recorded IAP richness in abandoned croplands and offers a baseline for future mechanistic, field-based, or monitoring-oriented research on invasion patterns in post-agricultural landscapes.

## 2. Materials and methods

### 2.1 Dataset

#### Data Synthesis and Taxonomic Standardization

To compile a national - scale overview of invasive alien plants (IAPs) reported in abandoned croplands, we conducted a literature-based data synthesis covering studies published between 1960 and 2024^31,32,34–59^. The dataset consists of occurrence records extracted from peer-reviewed articles that documented established IAP populations in abandoned cropland settings. Because the included studies varied in sampling design, spatial extent, and reporting detail, the resulting dataset reflects recorded occurrences rather than standardized ecological surveys. All species names were cross-referenced with international invasive plant databases to ensure taxonomic consistency. Invasion-risk categories were assigned following the framework of Yan et al. (2014) ^60^, which classifies species into five levels based on documented ecological impact and spread potential: malignant (Level 1), serious (Level 2), general (Level 3), local (Level 4), and species requiring observation (Level 5). As literature-based records typically emphasize established or widely recognized invaders, most species in the final dataset fell within Levels 1–4, whereas Level 5 species (requiring observation) were rarely reported. This synthesis approach prioritizes broad spatial coverage and taxonomic completeness over methodological uniformity, making the dataset suitable for exploratory macroecological analyses of recorded richness patterns.

#### Spatial Data Integration and Analytical Units

Because the compiled studies differed substantially in sampling protocols, geographic resolution, and survey periods, all explanatory variables were harmonized within a common analytical framework to enhance comparability across observations.

Two complementary richness metrics were used:

#### Cumulative provincial richness

Used only for descriptive mapping (Fig. 2), this metric represents the total number of unique IAP species recorded within each province across all studies. It reflects recorded richness, which may be influenced by variation in research intensity among provinces.

#### Study-level recorded richness

Used as the response variable in the GLMMs, this metric represents the number of IAP species reported in each individual study. Multiple observations may originate from the same province when different studies sampled distinct sites or periods.

Socio-economic variables (GDP, urbanization rate, population density, and HEIs) were matched to the corresponding province and sampling year of each study. Environmental variables (climate, soil, topography, and accessibility) were extracted from the geographic coordinates reported in each publication. Province was included as a random intercept to account for potential clustering among observations from the same administrative region.

Information on years since abandonment and successional stage was inconsistently reported across studies and therefore could not be standardized for inclusion. As a result, the dataset is best suited for examining broad-scale correlational patterns rather than temporal dynamics of post-abandonment succession.

#### Explanatory Variables

To explore how different socio-ecological contexts align with recorded IAP richness, we compiled 13 predictors grouped into environmental and socio-economic domains (Table 1).

1. Environmental variables

- Climatic factors: Four bioclimatic variables from WorldClim v2.1 were included: mean annual temperature (Tavg), maximum temperature (Tmax), minimum temperature (Tmin), and annual precipitation.
- Soil properties: Soil organic carbon (SOC) and total nitrogen (TN) were extracted from SoilGrids 2.0.
- Topographic factors: Elevation, slope, and aspect were derived from the SRTM v4.1 DEM.
- Environmental variables represent local site conditions based on study coordinates.
2. Socio-economic variables

- Anthropogenic context: Population density and urbanization rate were used as indicators of human activity.
- Economic development: GDP was included as a regional - scale contextual variable reflecting economic activity and infrastructure.
- Accessibility: Travel time to cities (Weiss et al., 2018) was used as a broad proxy for landscape connectivity.
- Sampling-effort proxy: The number of Higher Education Institutions (HEIs) in each province was included to approximate potential variation in research intensity. This variable does not directly measure sampling effort but provides a coarse indicator of where scientific activity may be more concentrated.

**Table 1.** Final set of selected variables in analytical models.

| Variable type | Variable sub-type | Variable description | Source | Spatial resolution |
| --- | --- | --- | --- | --- |
| Socio-economic | Economic Development | GDP (Gross Domestic Product) | National Bureau of Statistics (NBS) | Province-year |
|  | Urbanization | Urbanization rate | Regional Statistical Yearbooks | Province-year |
|  | Human Pressure | Human population density | Regional Statistical Yearbooks | Province-year |
|  | Accessibility | Global travel time to cities | Weiss et al., 2018 | Site coordinate |
|  | HEIs | Higher Education Institutions | Ministry of Education of the People's Republic of China | Province-year |
| Environmental | Climatic | Tavg (Mean Annual Temperature) | WorldClim v2.1 | Site coordinate |
|  |  | Tmax (Max. Temperature) | WorldClim v2.1 | Site coordinate |
|  |  | Tmin (Min. Temperature) | WorldClim v2.1 | Site coordinate |
|  |  | Precipitation | WorldClim v2.1 | Site coordinate |
|  | Soil | Soil Organic Carbon (SOC) | SoilGrids 2.0 (ISRIC) | Site coordinate |
|  |  | TN (Total Nitrogen) | SoilGrids 2.0 (ISRIC) | Site coordinate |
|  | Topographic | Elevation (m) | SRTM v4.1 (NASA/CGIAR-CSI) | Site coordinate |
|  |  | Slope (degree) | Derived from SRTM DEM | Site coordinate |
|  |  | Aspect (degree) | Derived from SRTM DEM | Site coordinate |

Socio-economic variables represent regional contextual conditions rather than local site characteristics.

#### Data limitations and analytical implications

Because richness values were derived from heterogeneous published studies rather than standardized field surveys, they may reflect differences in sampling extent, detection probability, and research focus. The inclusion of Province as a random effect and HEIs as a sampling-effort proxy helps mitigate, but cannot fully eliminate, these sources of variation. Consequently, the dataset is most appropriate for exploratory macroecological analyses aimed at identifying broad -scale correlational patterns rather than inferring causal relationships.

### 2.2 Statistical analysis

To explore broad - scale correlational patterns in recorded invasive alien plant (IAP) richness, we implemented a generalized linear mixed modelling (GLMM) framework. This approach is suitable for count - based response variables and for datasets in which multiple observations originate from the same administrative region^62^. All statistical analyses were conducted in R 4.3.0^63^.

We followed a four - stage analytical workflow designed to ensure transparency and comparability across heterogeneous literature-derived observations. First, we evaluated 14 candidate predictors for multicollinearity using variance inflation factors (VIF). A two-step screening procedure was applied: variables exhibiting extreme collinearity (e.g., Tmin and Tavg with VIF > 5) were removed to reduce redundancy, and all remaining predictors, including the sampling-effort proxy (HEIs), were retained only if their VIF values were below five ^64^. This procedure ensured that the final predictor set was sufficiently independent for exploratory modelling and that coefficient estimates were not unduly inflated by linear dependencies among variables.

Second, because study-level recorded richness is a discrete count variable, we fitted a Poisson GLMM with a log-link function using the *lme4* package ^65^. Continuous predictors were log - transformed where appropriate (e.g., GDP) and standardized (z - score transformation) to improve model convergence and facilitate comparison of effect sizes across variables measured on different scales. Although predictors were standardized, effect sizes were interpreted using incidence rate ratios (IRRs), obtained through exponential transformation of model coefficients (exp(β)), to provide an intuitive representation of multiplicative changes in expected recorded richness associated with a one-standard-deviation change in each predictor.

Third, Province was included as a random intercept to account for potential non - independence among observations originating from the same administrative region. This specification acknowledges that studies conducted within the same province may share unmeasured contextual characteristics, such as historical land-use patterns, reporting intensity, or regional research capacity, that could influence recorded richness. The random-effects structure therefore helps reduce pseudo-replication and allows fixed-effect estimates to reflect broad-scale correlational patterns rather than province-specific conditions.

Finally, model adequacy was evaluated using simulated residuals. Diagnostic checks indicated no evidence of overdispersion or zero inflation. Spatial autocorrelation was assessed using Global Moran’s I applied to simulated quantile residuals (DHARMa package) ^66^, with no significant spatial structure detected (p = 0.555). To avoid artificially inflating spatial dependence due to repeated coordinates across studies, residuals were aggregated by unique geographic locations prior to testing. Standardized effect sizes and marginal effects were visualized using the sjPlot package ^67^ to provide an interpretable overview of how each predictor aligned with variation in recorded richness.

Given the heterogeneous nature of literature-derived data and the limited number of study units, the GLMM results are interpreted as exploratory associations rather than causal relationships. The modelling framework is therefore intended to identify broad-scale correlational patterns that may inform future field-based or mechanistic research on invasive plant dynamics in post-agricultural landscapes.

## 3. Results

### 3.1. Composition of invasive plant species in abandoned croplands

Across all compiled studies, 57 invasive alien plant (IAP) species were recorded in abandoned croplands, representing 46 genera and 18 families. Asteraceae was the most frequently reported family (25 species; 43.9%), followed by Amaranthaceae (10 species) and Poaceae (5 species) (Fig. 1a).

**Fig. 1.**
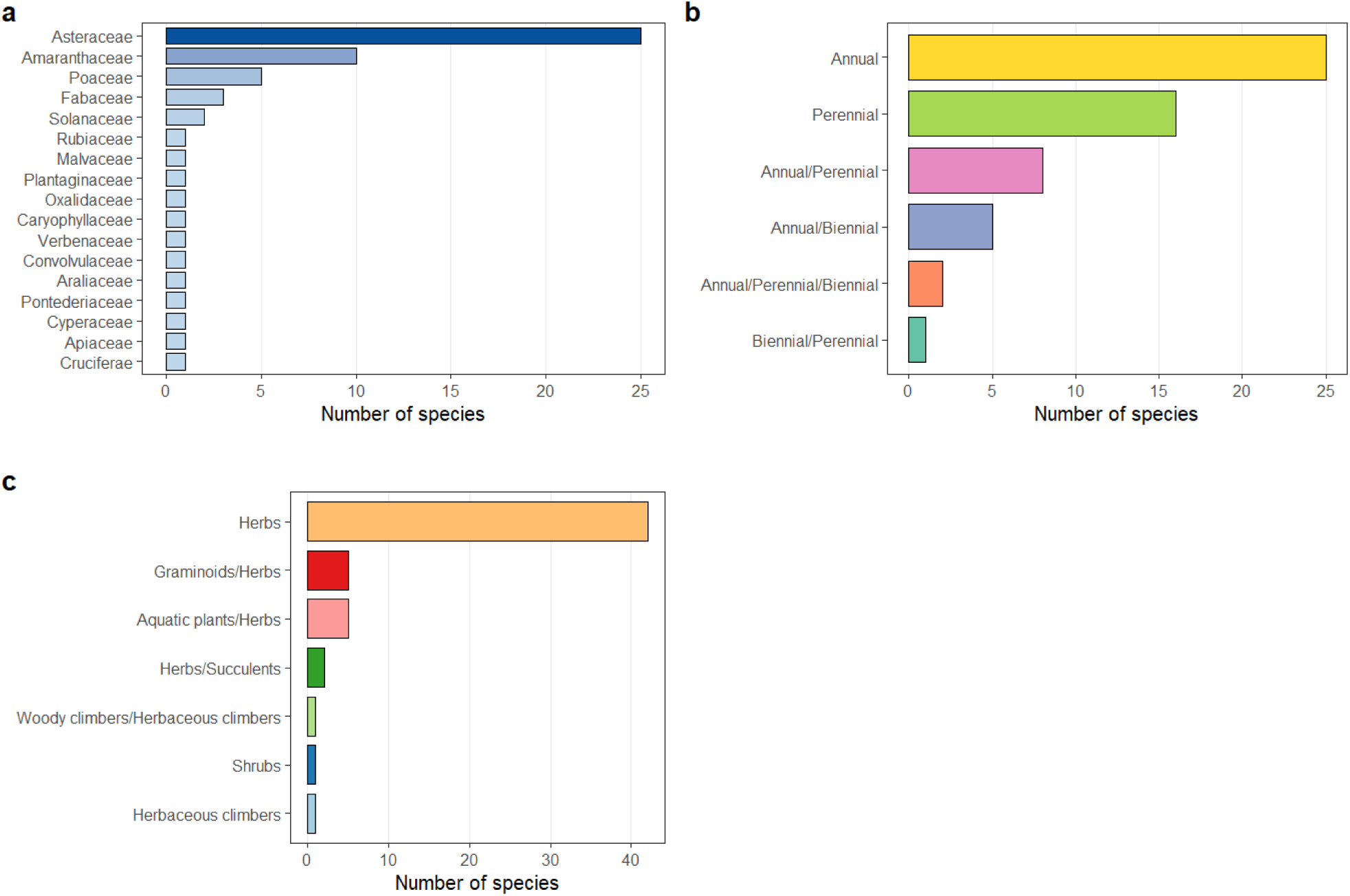
Taxonomic composition and functional trait structure of invasive alien plants (IAPs) recorded in abandoned croplands across China. (a) Family-level species richness; (b) life-history strategies (annual, biennial, and perennial species); (c) growth form composition (herbaceous plants, shrubs, succulents, and aquatic plants).

Most recorded species were herbaceous, reflecting a predominance of short-lived and disturbance-associated growth forms in the compiled literature (Fig. 1b). Annual species constituted the largest proportion, followed by perennials, whereas biennial and woody taxa were less commonly reported (Fig. 1c).

Based on the risk - classification framework of Yan et al. (2014), more than 70% of recorded species fell within Level 1 or Level 2 categories (Table 2). These patterns reflect the types of species most frequently documented in published studies of abandoned croplands rather than standardized assessments of ecological impact.

**Table 2.** Invasion risk levels of identified IAPs. Classification of 57 species into risk categories (Level 1 to Level 4) based on ecological impact and dispersal capacity as categorized by Yan et al. (2014).

| Invasion Risk Level | Category | No. of Species | Percentage (%) |
| --- | --- | --- | --- |
| Level 1 | Malignant invasion | 24 | 42.1 |
| Level 2 | Serious invasion | 18 | 31.6 |
| Level 3 | Local invasion | 3 | 5.3 |
| Level 4 | General invasion | 12 | 21.0 |
| Total |  | 57 | 100.0 |

### 3.2. Spatial pattern of recorded invasive plant richness

Recorded IAP richness exhibited marked spatial heterogeneity across China (Fig. 2). Higher cumulative provincial richness was observed in southeastern coastal provinces, with Guangdong reporting the largest number of unique species (30). Hainan (10 species) and Zhejiang (9 species) also showed relatively high cumulative richness.

**Fig. 2.**
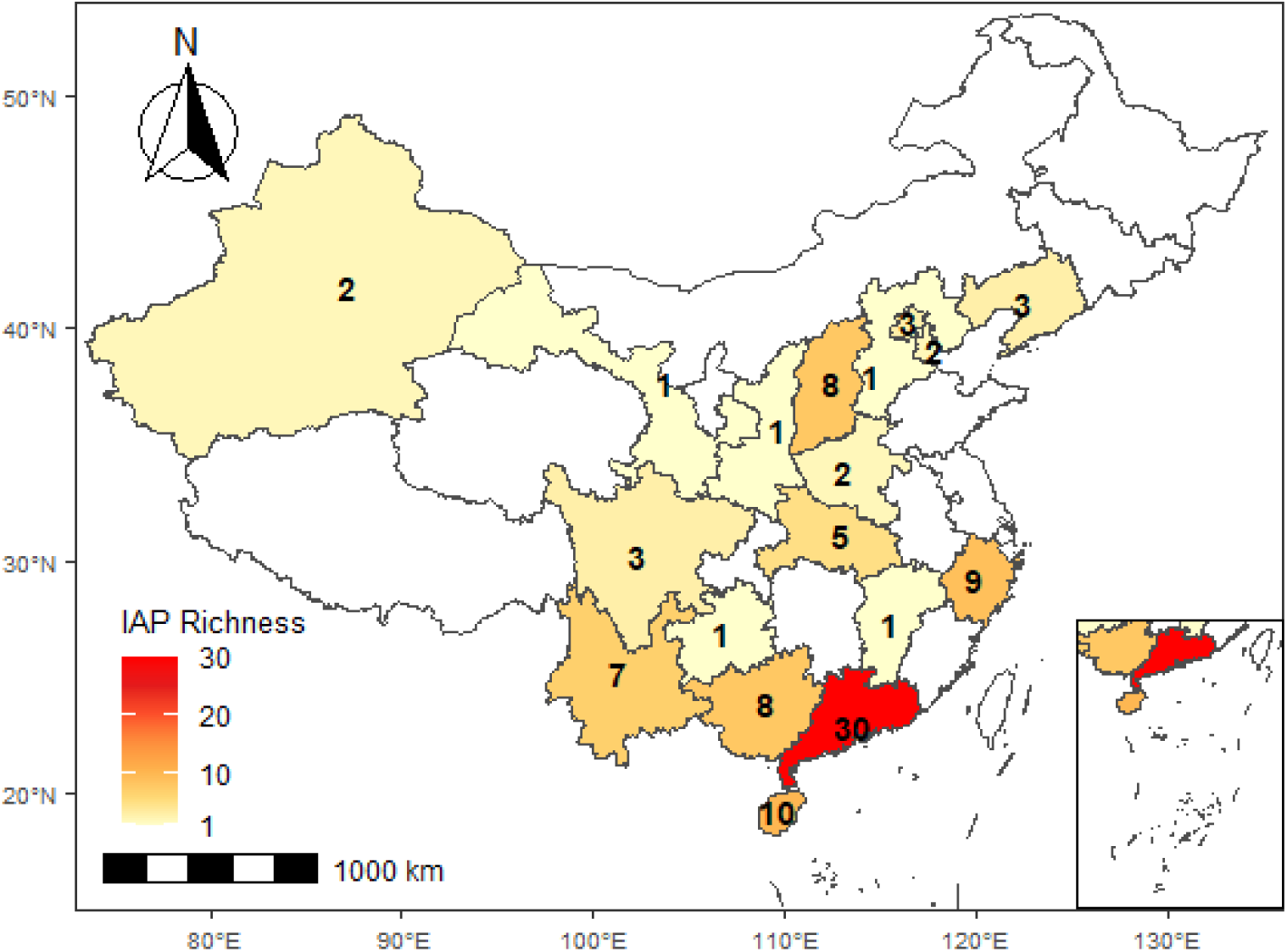
Cumulative provincial richness of invasive alien plants (IAPs) recorded in abandoned croplands across China. Richness values represent the total number of unique invasive species compiled from all literature-derived observations within each province.

In contrast, inland and northern provinces generally had fewer recorded species, with regions such as Sichuan, Beijing, and Tianjin reporting fewer than five species. These spatial patterns reflect variation in where species have been documented in the literature, which may be influenced by both ecological gradients and differences in research intensity across provinces.

### 3.3. Correlates of study-level recorded richness

After multicollinearity screening (VIF < 5; Table S1), socio-economic, climatic, soil, and topographic variables were retained for inclusion in the exploratory GLMM.

The model indicated that regional GDP (IRR = 1.67, p < 0.05) and mean annual temperature (IRR = 1.49, p < 0.05) were positively associated with study-level recorded richness (Fig. 3a). Other predictors, including elevation, slope, aspect, soil nitrogen, accessibility, and the sampling-effort proxy (HEIs), did not show detectable associations in the final model.

**Fig. 3.**
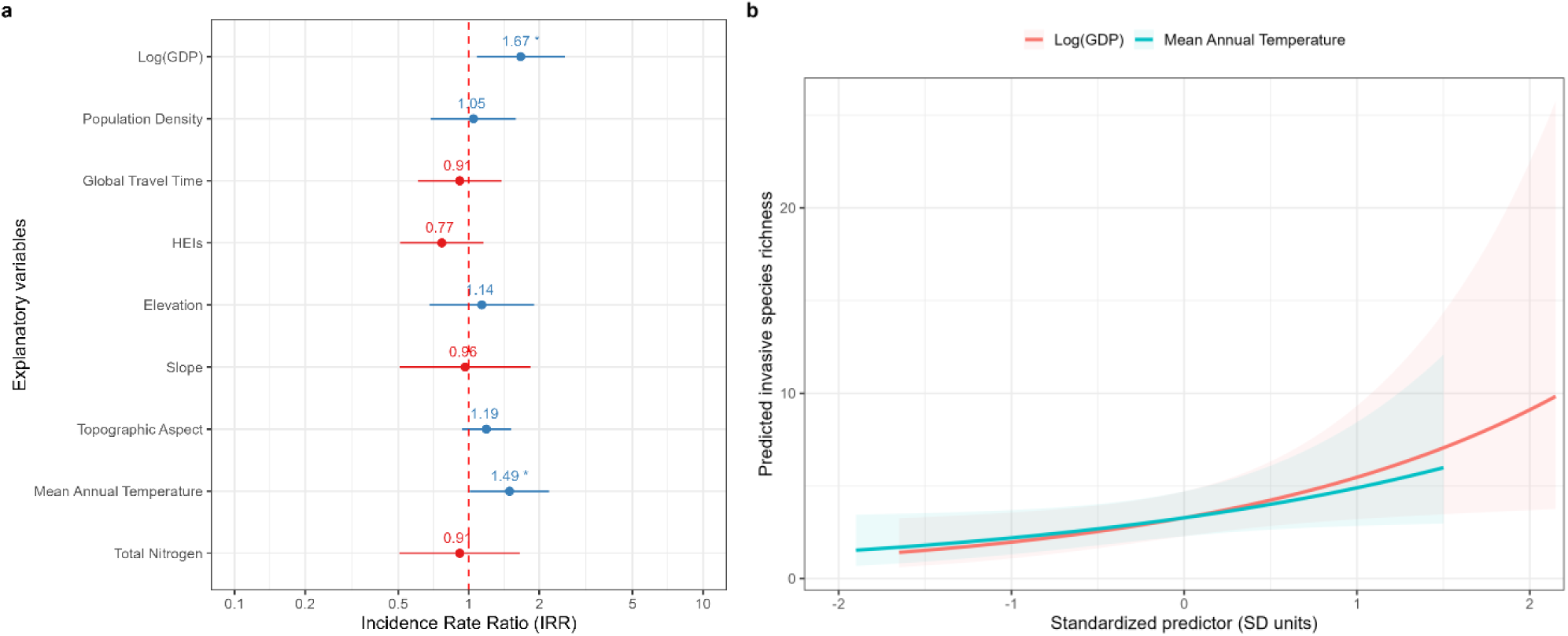
Results of the Generalized Linear Mixed Model (GLMM) for invasive alien plant richness. (a) Incidence rate ratios (IRRs) and 95% confidence intervals for all predictors retained after multicollinearity screening (VIF < 5) and included in the final GLMM; (b) marginal effects of significant predictors on predicted species richness. Asterisks indicate significance levels: *** p < 0.001, ** p < 0.01, * p < 0.05.

Marginal-effect plots showed gradual increases in predicted recorded richness across lower values of GDP and temperature, with steeper increases at higher values of both variables (Fig. 3b). These patterns represent statistical associations within the compiled dataset and do not imply causal relationships.

Model diagnostics indicated no evidence of overdispersion or zero inflation. Spatial autocorrelation tests based on simulated quantile residuals showed no significant spatial structure (Global Moran’s I, p = 0.555). Residuals were aggregated by unique coordinates prior to testing to avoid artificial clustering effects.

Interaction terms between key socio - economic and climatic variables were also examined, but no significant interactions were detected (p = 0.881).

## 4. Discussion

### 4.1 Abandoned croplands as socio - ecological contexts for recorded invasive plant assemblages

The compiled literature indicates that abandoned croplands in China are frequently associated with invasive alien plant (IAP) assemblages dominated by disturbance - adapted taxa. The predominance of Asteraceae, Amaranthaceae, and Poaceae reflects patterns commonly reported in studies of post - agricultural and human - modified environments, where short -lived, fast-growing species are more often documented. These families include many taxa that are widely recognized in the invasion literature, and their prominence in our synthesis likely reflects both their ecological characteristics and their higher detectability and reporting frequency in published studies.

The strong representation of herbaceous growth forms and annual life histories suggests that the IAPs recorded in abandoned croplands tend to align with traits commonly associated with early-successional or disturbance-linked vegetation. However, because the dataset is derived from heterogeneous studies with varying survey methods, these patterns should be interpreted as frequently reported assemblage characteristics rather than indicators of underlying ecological processes. The observed trait composition is therefore best viewed as a reflection of the species most commonly documented in the literature rather than a standardized assessment of trait filtering or successional dynamics.

More broadly, the assemblage patterns observed here are consistent with findings from other post-agricultural landscapes^20,68–71^, where species with high reproductive output. broad environmental tolerance, and rapid growth are often reported^72–74^. While such traits are frequently discussed in invasion ecology as being associated with successful establishment in disturbed environments, the present synthesis does not evaluate mechanisms directly. Instead, the results highlight the types of invasive species that have been most commonly recorded in abandoned croplands across China and provide a descriptive baseline for future studies that may investigate ecological processes in more detail.

### 4.2 Socio-ecological drivers of invasion richness across spatial gradients

The southeast–northwest gradient observed in recorded invasive plant richness aligns with broad spatial patterns frequently reported in studies of plant invasions in China ^20,68,69^ and elsewhere ^70,71^. Regions in southeastern China tend to have higher levels of economic development, denser transportation networks, and warmer climatic conditions, whereas inland and northwestern regions are characterized by lower human population density and stronger climatic constraints ^75–77^. These contextual differences may contribute to the geographic variation in where invasive alien plants have been documented, although the present synthesis does not evaluate underlying ecological or socio-economic mechanisms directly.

Within the exploratory GLMM, regional GDP and mean annual temperature were positively associated with study-level recorded richness. GDP may reflect a suite of regional characteristics, including economic activity, infrastructure, and human mobility, that are often discussed in the invasion literature as being linked to opportunities for species introductions and spread ^75–77^. In this study, GDP functions as a broad contextual indicator rather than a mechanistic driver, and its association with recorded richness should be interpreted as a correlational pattern within the compiled dataset.

Similarly, the positive association between mean annual temperature and recorded richness is consistent with the idea that warmer regions tend to support a wider range of alien plant species in observational datasets. Warmer climates are often associated with longer growing seasons and reduced thermal constraints, but the present analysis does not assess physiological processes or establishment dynamics. Instead, the observed pattern indicates that studies conducted in warmer regions have more frequently reported a greater number of invasive species.

In contrast, topographic variables, soil nitrogen, accessibility, and the sampling-effort proxy (HEIs) did not show detectable associations once GDP and temperature were included in the model. This may suggest that, at the national scale and within the limits of literature - derived data, regional socio - economic and climatic context aligns more strongly with variation in recorded richness than local environmental heterogeneity. However, the absence of detectable associations should not be interpreted as evidence of ecological irrelevance; rather, it reflects the constraints of the available data and the exploratory nature of the analysis.

Overall, the results highlight broad-scale correlational patterns between regional context and recorded invasive plant richness in abandoned croplands. These findings provide a descriptive foundation for future research that may incorporate standardized field surveys, finer-scale environmental measurements, or longitudinal data to more directly investigate ecological processes underlying invasion patterns in post-agricultural landscapes.

### 4.3 Abandoned croplands as transitional socio-ecological systems

Abandoned croplands in China can be viewed as transitional socio-ecological settings that retain characteristics of both agricultural and semi-natural landscapes. Although active cultivation ceases, many sites remain embedded within broader human-modified environments, including proximity to settlements, transportation corridors, and residual management structures ^81,82^. These contextual features may coincide with conditions under which invasive alien plants are more frequently documented in published studies, particularly where disturbance legacies such as altered soil structure or reduced vegetation cover persist ^13,78^.

Within this transitional context, the patterns observed in our synthesis suggest that recorded invasive plant richness aligns with both regional socio-economic conditions and broad climatic gradients. Warmer regions with higher levels of economic activity tended to have studies reporting a greater number of invasive species, whereas cooler or less densely populated regions generally had fewer recorded species. These associations reflect patterns within the available literature rather than underlying ecological processes, and the present dataset does not allow inference regarding mechanisms such as dispersal pathways, establishment dynamics, or competitive interactions.

Overall, abandoned croplands appear to function as heterogeneous socio -ecological contexts in which recorded invasive plant assemblages vary across regions. The descriptive patterns identified here provide a basis for future research that may incorporate standardized field surveys, abandonment histories, or spatially explicit ecological data to examine ecological processes more directly.

### 4.4 Implications for land-use transition governance

The descriptive patterns identified in this study have potential relevance for discussions surrounding land-use transitions in China, particularly in regions experiencing rapid socio - economic change ^83^. Current policy frameworks related to cropland abandonment primarily emphasize food security, agricultural productivity, and ecological restoration, while considerations related to invasive alien plants are less frequently integrated into land-use planning.

Our synthesis indicates that studies conducted in economically developed and climatically favorable regions have more frequently reported higher invasive plant richness. While these patterns do not imply causal relationships or predict future invasion dynamics, they suggest that regional socio - economic and climatic contexts may be relevant when considering how land - use transitions intersect with biological invasions. Integrating observational data on invasive species into broader land-use discussions may therefore support more comprehensive assessments of ecological outcomes associated with cropland abandonment.

More broadly, the findings highlight that ecological restoration and land-use transition processes can coincide with diverse ecological observations, including the presence of invasive species in some contexts. Continued monitoring and the incorporation of standardized ecological data would help clarify how different land-use trajectories relate to patterns of invasive plant occurrence across China’s rapidly changing landscapes.

### 4.5 Limitations and future directions

Several limitations of this study should be acknowledged. First, the dataset is derived from heterogeneous published studies that differ in sampling design, spatial resolution, survey effort, and reporting detail. As a result, recorded richness reflects the species documented in the literature rather than standardized assessments of invasive plant diversity. Variation in detection probability, plot size, survey timing, and taxonomic focus across studies may contribute to differences in recorded richness that cannot be fully accounted for in the present analysis.

Second, information on abandonment history, such as years since abandonment, prior land-use intensity, or management legacies, was inconsistently reported and therefore could not be incorporated into the analytical framework. These factors are frequently discussed in the invasion literature as relevant contextual variables, and their absence limits the ability to examine how temporal or historical dimensions align with recorded richness patterns.

Third, although the GLMM included a sampling - effort proxy (HEIs) and a random intercept for province, these approaches only partially address variation in research intensity and spatial clustering. Additional unmeasured factors, such as local monitoring capacity, publication bias, or regional research priorities, may also influence where invasive species have been documented.

Finally, the relatively small number of study units constrains the statistical power of multivariate modelling and limits the ability to explore interactions or nonlinear relationships among predictors. The results should therefore be interpreted as broad-scale correlational patterns rather than evidence of ecological processes or causal pathways.

Future work would benefit from standardized field surveys, consistent reporting of abandonment histories, and integration of finer - resolution environmental and socio - economic data. Longitudinal monitoring and spatially explicit modelling could also help clarify how invasion patterns evolve across post-agricultural landscapes. Despite these limitations, the present synthesis provides an initial descriptive baseline that may inform subsequent research on invasive alien plants in abandoned croplands.

## 5. Conclusion

This study provides an exploratory, national-scale synthesis of invasive alien plants (IAPs) recorded in abandoned croplands across China. By compiling heterogeneous literature-derived occurrence records and integrating them with socio-economic and environmental contextual variables, the analysis offers a descriptive overview of the species most frequently reported, the spatial distribution of recorded richness, and the broad-scale correlates associated with variation in study-level richness.

Across the compiled studies, recorded IAP assemblages were dominated by herbaceous and disturbance-associated taxa, with Asteraceae, Amaranthaceae, and Poaceae being the most frequently reported families. Spatially, higher cumulative provincial richness was documented in southeastern coastal regions, whereas inland and northern provinces generally showed lower recorded richness. These patterns reflect variation in where invasive species have been documented in the literature rather than standardized assessments of ecological distributions.

Within the exploratory GLMM, regional GDP and mean annual temperature were positively associated with study-level recorded richness, while other predictors showed no detectable associations. These findings represent correlational patterns within the available dataset and should not be interpreted as evidence of causal relationships or underlying ecological mechanisms. Instead, they highlight the types of regional contexts in which higher numbers of invasive species have been reported in published studies of abandoned croplands.

Given the heterogeneity of the underlying literature, the limited number of study units, and the absence of standardized field protocols, the results are best viewed as an initial macroecological overview. Future research incorporating systematic field surveys, finer-resolution environmental data, and temporal information on post - abandonment succession would be valuable for assessing ecological processes more directly. Nonetheless, the present synthesis provides a baseline reference for understanding broad - scale patterns in recorded IAP richness and may help guide subsequent monitoring and research efforts in post-agricultural landscapes.

## Acknowledgments

This research was supported by the Postgraduate Research Scholarship (PGRS) at Xi’an Jiaotong-Liverpool University (No. PGRS2312003). This work was conducted as part of a doctoral dissertation at the University of Liverpool, in collaboration with Xi’an Jiaotong-Liverpool University, and constitutes a core component of the first author’s PhD thesis.

During the preparation of this manuscript, ChatGPT (OpenAI) was used solely to assist with language editing and refinement. All scientific content, analyses, interpretations, and conclusions were developed, verified, and approved by the authors, who take full responsibility for the content of this manuscript.

## 7. Supporting information

**Table S1.**
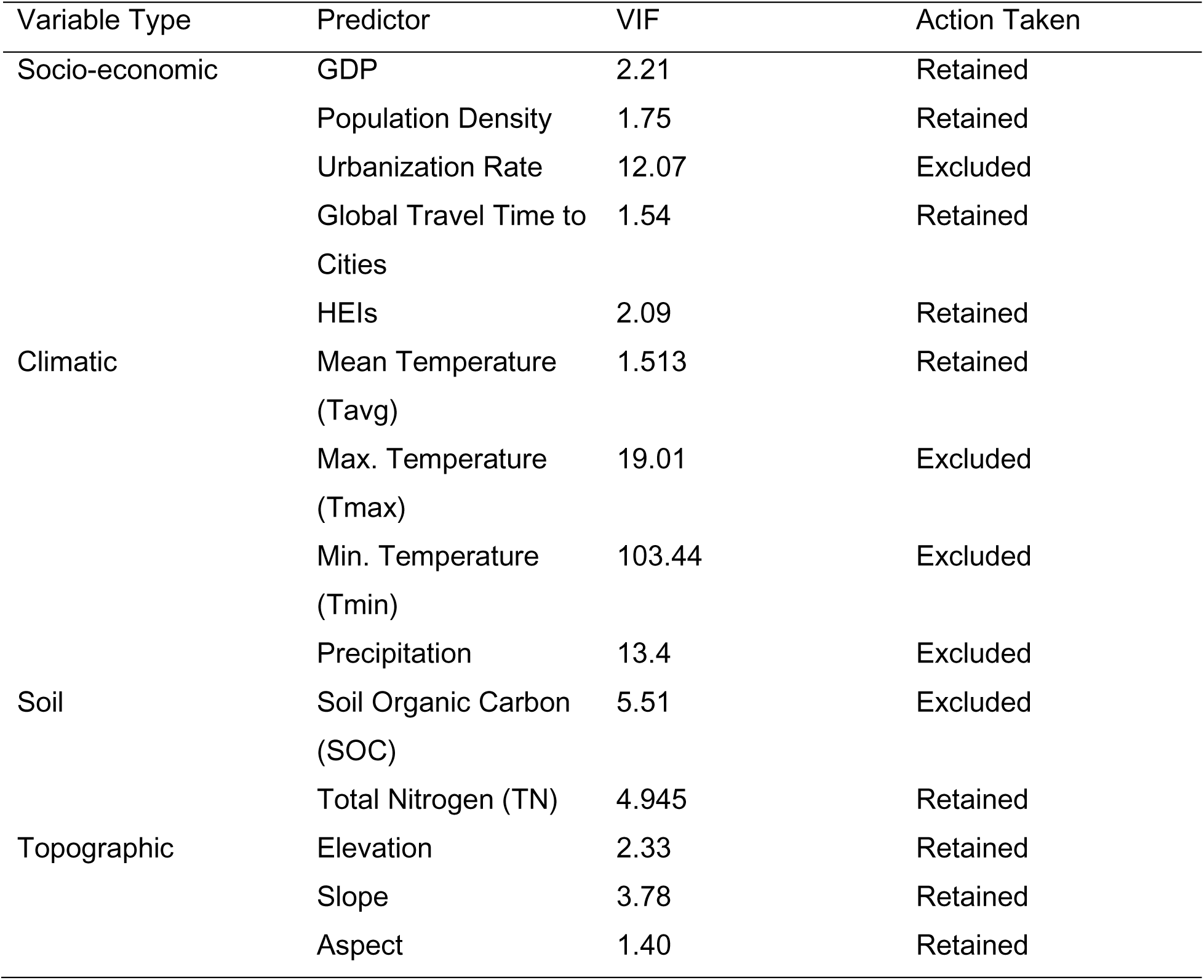
Multicollinearity diagnostics for the 13 initial predictor variables.

